# Streamlined isolation of the n-terminome via phosphonate tagging and coordination-based depletion

**DOI:** 10.1101/2025.05.30.657046

**Authors:** Piero Giansanti, Adnan Fojnica, Andreas Pichlmair

## Abstract

Protein degradation is critical for regulating cellular functions. Therefore, there is considerable interest in developing effective proteome-wide strategies to identify protease cleavage products to enhance our understanding of proteolytic pathways and their perturbation in diseases. Here, we present a streamlined N-termini proteome analysis leveraging N-Hydroxysuccinimide (NHS) ester chemistry. At the protein level, N-terminal amines (naturally occurring protein N-termini, lysines, or protease-generated N-termini) are blocked directly in the cell lysate, and tryptic digestion is performed straight after, without the need for any buffer exchange. The internal tryptic peptides are tagged with a phosphonate moiety and subsequently depleted via metal-based coordination, keeping exclusively N-blocked terminal peptides for analysis by LC-MS/MS. We demonstrate the applicability of this approach by monitoring proteolytic events induced by intrinsic apoptotic pathway. Our approach identified more than 690 cleaved proteins in response to Staurosporine-induced apoptosis, including many previously unknown substrates and cleavage sites. The presented approach is, therefore, a straightforward and robust method for mass spectrometry-based identification of caspase-generated cleavage products and extendible to a wide range of other proteolytic cleavage events.

## INTRODUCTION

Proteolysis by endogenous proteases is a fundamental regulatory mechanism that impacts nearly every aspect of cellular function, from signal transduction to differentiation, protein turnover, and immune responses^1–3^. Many physiological and pathological processes, including cancer progression, inflammation, and viral infections, are also driven by protease activity^4–6^. Understanding proteolytic events is, therefore, crucial for deciphering biological processes and disease mechanisms, and it also facilitates the development of targeted therapeutics and biomarkers. In fact, by characterizing protease-substrate interactions, novel regulatory mechanisms can be uncovered, and potential drug targets for therapeutic intervention can be identified^7–9^.

Despite its importance, systematically mapping protease substrates and cleavage sites remains a challenging task due to the transitory and dynamic nature of proteolytic events^10^. Mass spectrometry-based proteomics has played a critical role in systematically characterizing proteolysis at a global scale^11–15^. Cravatt and coworkers pioneered a proteomic approach that enabled proteome-wide identification of protease substrates^16^, later elegantly extended to a quantitative level using SILAC^17,18^. These methods enable the identification of proteolytic events by detecting shifts in protein migration on SDS-PAGE, providing insights into protease activity and substrate specificity. However, they are effective when proteolytic events cause a substantial shift in molecular weight, and are generally limited in precisely mapping cleavage sites, in detecting low-abundance cleavage products, and in scaling for high-throughput analysis due to limited multiplexing and extensive MS analysis required. Therefore, positional proteomics^19^, and in particular, N-terminomics has emerged as a highly effective alternative for the in-depth and high-throughput characterization of proteolytic cleavage events^20^, which addresses some of the limitations of the aforementioned methods. The rationale behind N-terminomics is that proteolysis generates new N-termini, which can be selectively exploited to determine precise cleavage sites. Techniques such as TAILS^21^, and HUNTER^22^ leveraged this principle to scavenge for protease-generated N-termini by taking advantage of the unique chemical structure and reactivity of the primary amines, enabling the direct identification of cleavage sites with high sensitivity.

These approaches follow a general workflow consisting of five main steps: : i) protein extraction and denaturation to improve the accessibility of amino groups, ii) blocking/labeling of both protein N-termini (canonical or protease-generated) and lysine side chains, leveraging their readily reactive α- and ε-amines, iii) proteome digestion, iv) chemical tagging of internal peptides generated upon digestion containing newly exposed α-amines, and v) selective depletion of internal peptides making use of the introduced tag.

Despite their efficacy, these methods have several drawbacks that limit their applicability for high-throughput analysis: the experimental workflow is lengthy and tedious, requiring two rounds of proteome cleanup, either through standard organic solvent-induced precipitation alone or in combination with capture onto hydroxylated magnetic beads^23,24^ to remove interfering compounds from the protein lysate, and before digestion. This not only increases time and cost but also introduces variability due to potential sample loss, especially since small cleavage products do not precipitate as efficiently as larger proteins^22,25,26^. Additionally, despite cleanup, blocking/labeling reagents are typically used in high molar excess, with protein-to-reagent ratios ranging from 1:4 to 1:16, for TMT10 or TMTpro^23,27,28^. Lastly, the negative selection of the N-terminome relies on a Schiff’s base reaction using cyanoborohydride, a highly toxic reagent that also produces hazardous byproducts, necessitating careful handling.

In view of these challenges, we have developed SIMPATICO, a streamlined isolation of the n-terminome via phosphonate tagging and coordination-based depletion. The method is built on the PTAG approach^29^, but it radically minimizes processing steps and eliminates the need for toxic reagents. It relies exclusively on the well-established and widely used NHS esters chemistry, employing inexpensive reagents and standard laboratory equipment, making it easily accessible for any proteomics laboratory.

We demonstrate that SIMPATICO is a straightforward and efficient strategy, enabling the identification of more than 900 cleavage events in response to Staurosporine-induced apoptosis, and it can seamlessly integrate with multiplexing and label-free quantitative analysis.

## EXPERIMENTAL PROCEDURES

### Cell Culture, Stimulation and Lysate Preparation

HeLa S3 cells were cultured in DMEM medium (Sigma-Aldrich), while Jurkat cells were maintained in RPMI 1640 (Sigma-Aldrich), both supplemented with 10% fetal calf serum (GE Healthcare) and antibiotics (100 U/ml penicillin, 100 µg/ml streptomycin, Sigma-Aldrich). Cells were incubated at 37 °C in a humidified atmosphere (95% humidity, 5% CO₂) and routinely tested for *Mycoplasma* contamination.

Jurkat cells were seeded at 1 × 10^6^ cells/mL before apoptosis induction. Staurosporine (STS, 1 µM, MedChemExpress) was added, and the cells were incubated for 6 h at 37 °C. HeLa and STS-treated Jurkat cells were then harvested, washed three times with PBS (Sigma-Aldrich), snap-frozen in liquid nitrogen, and stored at −80°C until further use. Pellets were thawed on ice and cells were lysed in lysis buffer containing 2 or 4% SDC (w/v, Sigma) and 200 mM HEPPS (Sigma), pH 8.5, by heating them for 5 min at 95 °C under shaking at 1500 rpm. The lysates were then sonicated with a Bioruptor Pico (Diagenode) for 10 cycles ON/OFF of 30 s. Protein concentration was estimated using a BCA assay (Thermo Fisher). To reduce and alkylate proteins, samples were incubated for 10 min at 45 °C with 10 mM TCEP (Thermo Scientific) and 40 mM CAA (Sigma).

### Primary Amines Blocking

The protocol for NHS ester-mediated blocking of primary amines was adapted from Zecha et al.^30^.

For initial method optimization, individual channels of TMT10plex reagent (Thermo Scientific) were brought to RT and dissolved in anhydrous ACN under Argon gas. Varying amounts of reagents were mixed with varying amounts of cell lysate, maintaining a final ACN concentration of 20% (v/v), and incubated for 1 hour at 20 °C in a thermoshaker with shaking at 500 rpm. Respective volumes and concentrations are specified in the results section. Afterward, ABC (Sigma) was added to quench the reaction to achieve a final concentration of 50 mM, obtaining a final concentration of 0.24% SDC (w/v) and 6% ACN (v/v) for the lysate containing 1% SDC (v/v), and of 0.4% SDC (w/v) and 5% ACN (v/v) for the lysate containing 2% SDC (v/v).

For the multiplexing experiments, 0.6 mg of TMT10plex reagents (Thermo Scientific) were dissolved in anhydrous ACN, and added to the samples (200 μg in 125 μL, 1% SDC), ensuring a final ACN concentration of 20% by volume. The blocking reaction proceeded as described above. ABC was added to quench the reaction to achieve a final concentration of 50 mM, obtaining a final concentration of 0.24% SDC (w/v) and 6% ACN (v/v). Quenching was performed for 30 minutes at 25 °C and 500 rpm, after which the labeled samples were combined.

For the label-free experiments, 2,5-dioxopyrrolidin-1-yl 2-methyl-5,8,11-trioxa-2- azatetradecan-14-oate (DMA-PEG3-NHS) was custom-synthesized by JenKem Technology upon request and used as blocking reagent for primary amine. Each sample received 0.67 mg of reagent dissolved in dry ACN while maintaining a final ACN concentration of 20% (v/v). Blocking and quenching were performed as described above.

### Protein Digestion and Cleanup

Digestion was performed by adding trypsin (Thermo Scientific) at a 1:50 enzyme-to-substrate ratio followed by incubation overnight at 37 °C with continuous shaking at 700 rpm. Afterward, water was added to reduce ACN concentration to 3.8% (v/v), and digestion was stopped by acidification with neat FA to a final concentration of 5% (v/v). SDC and other insoluble matter were removed via centrifugation at 4 °C for 10 min.

Solid-phase extraction cartridges were used for desalting. For experiments involving TMT-multiplexing, a 200 mg, 3cc C18 SepPak column (Waters) was used, whereas for label-free experiments, a 50 mg, 1cc tC18 SepPak column (Waters) was utilized for each sample. The C18 sorbent material was conditioned by washing with pure ACN, followed by 80% ACN/0.1% FA (v/v), and then re-equilibrated twice with 0.1% FA (v/v). After loading the sample onto the columns, they were washed once with 0.1% FA (v/v). The peptides were then eluted using 80% ACN/0.1% FA (v/v). The washing and re-equilibration volumes for the 3cc column were 2 mL, while for the 1cc columns, 1 mL was used. For the elution step, 1 mL or 600 μL was used depending on the column size. The eluates were then dried by vacuum centrifugation and stored at −80 °C until further use.

### Preparation of Phosphonate-tagged Internal Peptides

For initial method optimization, HeLa tryptic digest was resuspended in 200 mM HEPPS, pH 8.5 and mixed with varying amounts of 1-(6-Phosphonohexanoyloxy)-2,5- pyrrolidinedione (NHS-PnTAG, MedChemExpress, HY-28043), maintaining a final ACN concentration of 20% (v/v), and incubated for 1 hour at 20 °C in a thermoshaker with shaking at 500 rpm. Respective volumes and concentrations are specified in the results section.

For the multiplexing and label-free experiments, desalted peptides were reconstituted in 200 mM HEPPS (pH 8.5) to a concentration of 10 μg/μL and incubated for 10 min at 20 °C while shaking at 500 rpm. The free amino groups of the internal tryptic peptides were phosphonate-tagged by adding NHS-PnTAG at a final concentration of 13.7 mM in 20% ACN. The reaction was incubated for 1 hour at 20 °C and 500 rpm. To quench the tagging reaction, 100 μL of a 75 mM ABC solution was added to the mixture and incubated for 30 min at 25 °C at 500 rpm. Finally, water was added to reduce the ACN concentration to 4% (v/v) and samples were acidified with FA to 5% (v/v). The peptides were desalted as previously described.

### High-pH Reversed Phase Fractionation

TMT-labeled and phosphonate-tagged peptides were subjected to offline basic pH reversed-phase fractionation. Peptides were reconstituted in 25 mM ammonium bicarbonate at pH 8.5 and loaded onto an XBridge BEH C18 column (Waters; 3.5 μm, 4.6 mm × 250 mm) connected to a Dionex Ultimate 3000 HPLC system (Thermo Scientific). Peptides were loaded for 3 min at 100% buffer A (95% H_2_O, 5% ACN, 2.5 mM ABC, pH 8.5), and separated at a flow rate of 1 mL/min using a 42.5-min linear gradient from 7% to 45% of buffer B (5% H_2_O, 95% ACN, 2.5 mM ABC, pH 8.5), followed by a 3.5-min washing with 85% B. A total of 96 fractions were collected, which were subsequently pooled into 24 fractions, acidified with FA to a final concentration of 0.5%, and dried by vacuum centrifugation at 45 °C.

### Depletion of Phosphonate-tagged Peptides

Phosphonate-tagged and fractionated peptides were depleted using 5 μL Fe(III)-NTA cartridges in an automated fashion on the AssayMAP Bravo Platform (Agilent Technologies), adapting a standard enrichment protocol^31^. In detail, cartridges were primed at a flow rate of 300 μL/min with 150 μL of priming buffer (0.1% TFA in ACN) and equilibrated at 10 μL/min with 150 μL of loading buffer (0.1% TFA in 80% ACN). Subsequently, peptides, dissolved in 200 μL of loading buffer, were loaded at 5 μL/min onto the cartridge, and the flow through was collected into a separate plate. Columns were washed with 150 μL of loading buffer at a flow rate of 10 μL/min, and the flow through was collected and combined with the previous one. Finally, bound peptides were discarded with 60 μL of 1% NH_4_OH (v/v). The depleted fractions were subsequently pooled into 12 fractions.

For label-free experiment, depletion was performed via solid-phase extraction using 30 mg, 1cc zirconia-based HybridSPE cartridges (Sigma). The columns were first conditioned with 500 μL of loading buffer (1% FA in 80% ACN). Afterward, samples were loaded in 250 μL of loading buffer, and the flow throughs were collected. Cartridges were sequentially washed twice with 400 μL of 1% FA in 50% ACN and 400 μL of 1% FA in 10% ACN. During all washing steps the flow through were collected, combined with the previous ones, and then dried by vacuum centrifugation at 45°C. Finally, depleted peptides were desalted using SDB-RPS StageTips (Empore) via centrifugation at 100- 500 x *g* for 2-5 min. To this end, peptides were reconstituted in 1% TFA and transferred to StageTips. After loading the samples, the StageTips were washed sequentially with 100 μL of 1 % TFA in IPA, 10 mM NH_4_FA in 50%ACN/5% FA, and 0.2% TFA. Peptides were eluted using 150 μL of 1% NH_4_OH in 80% ACN. The SDB-RPS StageTips cleanup procedure was also used to desalt samples of the titration experiments.

All depleted samples were dried down at 45 °C and stored in −20 °C until LC-MS/MS analysis.

### Nanoflow LC-MS/MS

The analysis of N-terminal peptides (depletion flow-through) was performed either on a Dionex Ultimate 3000 UHPLC+ coupled to an Orbitrap Eclipse mass spectrometer or on a Vanquish Neo UHPLC system coupled to an Orbitrap Exploris 480 mass spectrometer (all Thermo Fisher).

The Dionex system was equipped with a 2 cm trap column (75 μm i.d., packed in-house with 5 μm Reprosil C18 beads; Dr. Maisch) and a 45 cm analytical column (75 μm i.d., packed in-house with 1.9 μm ReproSil-Pur C18-AQ beads; Dr. Maisch). Trapping and washing were performed at 5 μL/min for 10 min with solvent A (0.1% FA in water). Subsequently, peptides were transferred to the analytical column at 300 nL/min in a total analysis time of 90 min with a gradient of 8%-34% (v/v) solvent B (0.1% FA, 5% DMSO in ACN) in 80 min. The mass spectrometer was operated in a data-dependent MS3 mode. Every 3 s, a MS1 survey scan was recorded from 360 to 1500 m/z with a resolution of 60k in the Orbitrap. The MS1 AGC target was set at 100%, and the maxIT was set to 50 ms. Precursors were targeted for HCD fragmentation with an NCE of 34% if the charge was between 2 and 6, and the intensity exceeded 2e4. The quadrupole isolation window was set to 0.7 Th. The MS2 spectra were acquired in the Orbitrap with a resolution of 15K, an AGC target at 100%, and maxIT of 22 ms. Precursors that have been targeted for fragmentation were excluded for 90 s for all possible charge stages. TMT reporter ions were measured in a consecutive MS3 scan based on the previous MS2 scan. Thus, a new batch of precursor ions was isolated with a quadrupole isolation window of 1.2 Th. The isolated precursor was then HCD-fragmented identically to the previous MS2 scan. The top 10 fragment ions of the MS2 scans were isolated in the ion trap in parallel via synchronous precursor selection, and HCD fragmented with an NCE of 55%. The MS3 spectrum was acquired with 30k resolution from 100 to 500 m/z in the Orbitrap, and the phase-constrained spectrum deconvolution method (PhiSDM) was enabled for the TMT reporter ions. The MS3 AGC target was set at 500%, and the maxIT was set to 54 ms.

The Vanquish Neo system was equipped with a 5 cm trap column and a 40 cm analytical column, both packed in-house as described above. Sample loading was performed using 0.1% FA at a flow rate of 5 μL/min, then peptides were transferred to the analytical column and separated at 300 nl/min using an 81-min linear gradient 4% to 25% of solvent B (0.1% FA, 3% DMSO in ACN) and solvent A (0.1% FA, 3% DMSO in water). The total analysis time was 90 min. The Orbitrap Exploris 480 was operated in data dependent and positive ionization mode. MS1 spectra were recorded in the Orbitrap from 360 to 1350 m/z at a resolution of 60K, with an AGC target value at 100%, and maxIT of 50 ms. The top 25 precursor ions in the MS1 scans with a charge between 2 and 6 were isolated via the quadrupole with an isolation window of 0.7 Th, and HCD-fragmented with a NCE of 36%. MS2 spectra were recorded in the Orbitrap at 15K resolution, with an MS2 AGC target value at 200%, and maxIT set to 22 ms. The PhiSDM option was enabled, and dynamic exclusion was set to 40 s.

Labeling and tagging efficiency samples were acquired on the Vanquish-Exploris system, with the following modifications: MS1 spectra were recorded from 360 to 1300 m/z with an AGC target value at 300%, the top 15 precursor ions in the MS1 were subjected to HCD fragmentation with an NCE of 30%, quadrupole isolation window was set to 1.3 Th, and MS2 spectra were recorded at 15K resolution with an AGC of 200% and maxIT set to 22 ms, Dynamic exclusion was 30 s.

Label-free samples were acquired on the Vanquish-Exploris system, with the following modifications: MS1 spectra were recorded from 400 to 1500 m/z with an AGC target value at 300%, the top 15 precursor ions in the MS1 were subjected to HCD fragmentation with a stepped NCE of 35 ± 5%, quadrupole isolation window was set to 1.3 Th, and MS2 spectra were recorded at either 15K of 30K resolution with a maxIT set to 22 or 54 ms, respectively. Dynamic exclusion was 30 s.

### Data Analysis

Raw mass spectrometry data were processed with the FragPipe^32^ software (version 22.0). Spectra were searched against the UniProtKB database (Human, UP000005640, 104,726 entries including isoforms, downloaded on 10.2023). Enzyme specificity was set to semi-trypsin_r with free N-terminus, allowing for two missed cleavages, and the search included cysteine carbamidomethylation as a fixed modification and methionine oxidation as a variable modification. TMT (+229.16293) or DMA-PEG3ylation (+231.14706) were fixed modifications on peptide n-terminus and lysine.

Separate searches were conducted to evaluate labeling efficiency. Thus, TMT (+229.16293) or DMA-PEG3ylation (+231.14706) was specified as a variable modification on primary amines (lysine and protein n-terminus) or as a fixed modification on primary amines and, additionally, as a variable modification on serine, threonine, and tyrosine. Enzyme specificity was set to strict_trypsin, and up to three or two missed cleavages were allowed, for labeling and overlabeling search, respectively. For tagging efficiency, 6-Phosphonohexanoylation (+178.03949) was specified as a variable modification on lysine and peptide n-terminus or as a fixed modification on primary amines and as a variable modification on serine, threonine, and tyrosine.

For TMT multiplexing data, peptide-to-spectrum matches (PSM) rescoring via MSBoster^33^ was enabled, using predictions from Prosit via the Koina service^34^. Finally, identifications were adjusted to 1% false discovery rate (FDR) with percolator^35^ and Philosopher^36^, integrated into FragPipe.

Data analysis was performed with the Perseus^37^ software (version 2.1.3.0). Peptide identifications were filtered to remove contaminants before performing data normalization of the log2-transformed peptide intensity values by median centering, as implemented in Perseus. For statistical analysis, only peptides quantified in at least three biological replicates in at least one experimental group were retained. Missing values were imputed from a normal distribution in Perseus using default parameters, but only for peptides that did not already meet this threshold (3 valid values) in both groups. Differential analysis (STS vs. Mock) was conducted using Welch’s t-test with a permutation-based FDR threshold of 5% to account for multiple testing.

## RESULTS AND DISCUSSION

### Workflow Optimization

We initially sought to evaluate the feasibility of performing primary amine blocking directly on the proteome lysate, immediately followed by digestion, without requiring any cleanup step. SDC has emerged as a widely used detergent in proteomics by virtue of enhancing trypsin activity and being easy to remove prior to LC-MS analysis^38^. Given that this detergent lacks a primary amine group and is soluble in both polar and non-polar solvents, we reasoned that it could be used effectively for cell lysis and in a one-pot blocking/digestion procedure, where the TMT reaction and digestion share a common buffer.

To test this hypothesis, a protein and TMT titration experiment was performed in triplicates using a fixed protein amount of 100 μg in varying volumes of lysis buffer (1 or 2% SDC, 200 mM HEPPS, pH 8.5) ranging from 50 to 75 μL, and different amounts of TMT (200 and 400 μg) in 20% ACN. The reaction was stopped by adding ABC to a final concentration of 50 mM, also lowering the SDC (0.24%) and ACN (6%) concentrations for direct trypsin digestion. Samples were analyzed using single-shot LC-MS/MS runs (Supplementary Table S1). Across the entire range of tested protein and TMT quantities, at least 98% of the detected peptides bearing labelable amines (protein N-termini and lysine side chains) were fully labeled (Figure 1A). As expected, the higher TMT-to-protein ratio resulted in higher labeling efficiency; however, the magnitude of this difference is negligible. Consequently, few unlabeled or partially labeled peptides were observed. Interestingly, the corresponding over-labeling analysis revealed that on average, just ∼6% of the detected peptides contained at least one TMT-labeled serine, threonine, or tyrosine residue (Figure 1B), with minimal difference across the various TMT concentrations.

**Figure 1.**
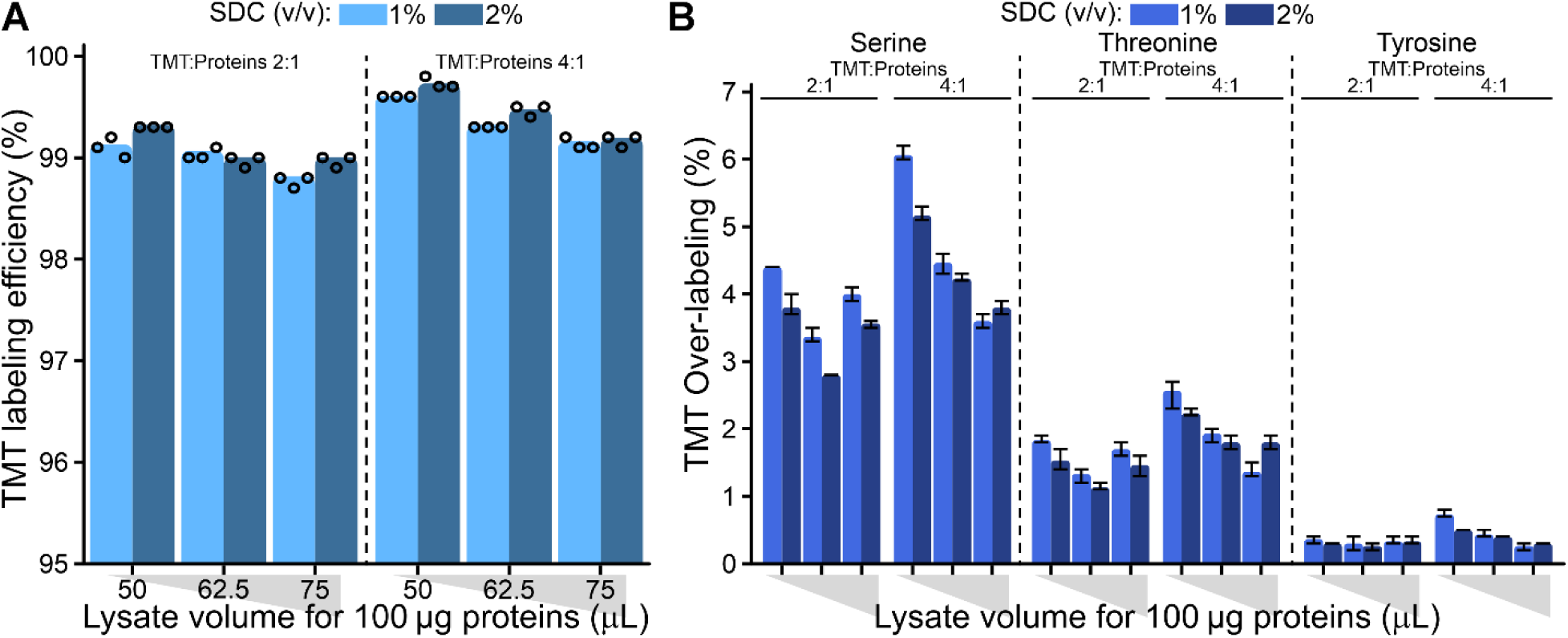
Evaluation of protein-level TMT labeling protocols directly on the lysate. (A) Bar plot of the percentage of peptide intensity from labeled peptides at lysines and protein N- termini (labeling efficiency) across different reaction volumes and amounts of TMT10 reagent. (B) Bar plot of the percentage of peptide intensity attributable to over-labeled, O- acylated peptides by ester type from the experiment displayed in (A). For all plots, bars are drawn at the mean value (n = 3), dots indicate individual replicates, and error bars indicate the minimum and maximum values.

Taken together, this indicates that blocking of primary amine directly on the cell lysate is possible, greatly facilitating the sample preparation and sidestepping variable sample losses due to multiple cleanups. We note that the abundance of primary amines varies significantly depending on the sample type, including contributions from metabolites, lipids, free amino acids, etc.; hence, to ensure complete labeling (>99%), while minimizing overlabeling, we identify 3:1 as a suitable TMT-to-protein ratio for subsequent experiments.

### NHS Ester-Mediated Phosphonate Tagging of Internal Peptides

Removal of internal peptides is essential in N-terminomics to specifically identify peptides originating from the N-terminus of proteins or newly exposed N-termini after proteolysis. This is typically achieved through a Schiff base reaction, where the α-amines of internal peptides react with a reagent that alters their hydrophobicity or molecular weight^39^, allowing them to be captured and removed. While the reaction is selective, it generally requires several hours to ensure complete derivatization. Additionally, it requires careful handling due to the potential hazards of the reagents or byproducts involved.

Here, we adapted a strategy based on crosslinking peptides with a phosphonate moiety using NHS chemistry^40^. By translating this approach, we tagged internal peptides with a phosphonate group using an inexpensive and fast-reacting NHS ester. Depletion is then seamlessly performed using metal-based coordination methods that are broadly used for phospho-peptide enrichment^41,42^.

To evaluate the efficiency of our tagging strategy, we used a HeLa tryptic digest and performed again a titration experiment using a fixed amount of peptides and increasing concentration of NHS-PnTAG, resulting in tag-to-peptide ratios of 1:4 up to 1:2 (Figure 2A and Supplementary Table S2). Single-shot LC-MS/MS analysis revealed nearly complete derivatization at a 1:2 ratio, with minimal over-tagging on serine, threonine, or tyrosine residue (Figure 2B). To rule out the formation of other unexpected products due to minor impurities in the NHS-PnTAG reagent (purity > 95% according to the manufacturer), we used the open search functionality of MSFragger to detect any new modifications on the peptides (Supplementary Table S2). Although we identified over 40 potential mass shifts, they only accounted for ∼15% of the identified peptides, and the majority of these were also present in the control experiments (Figure 2C).

**Figure 2.**
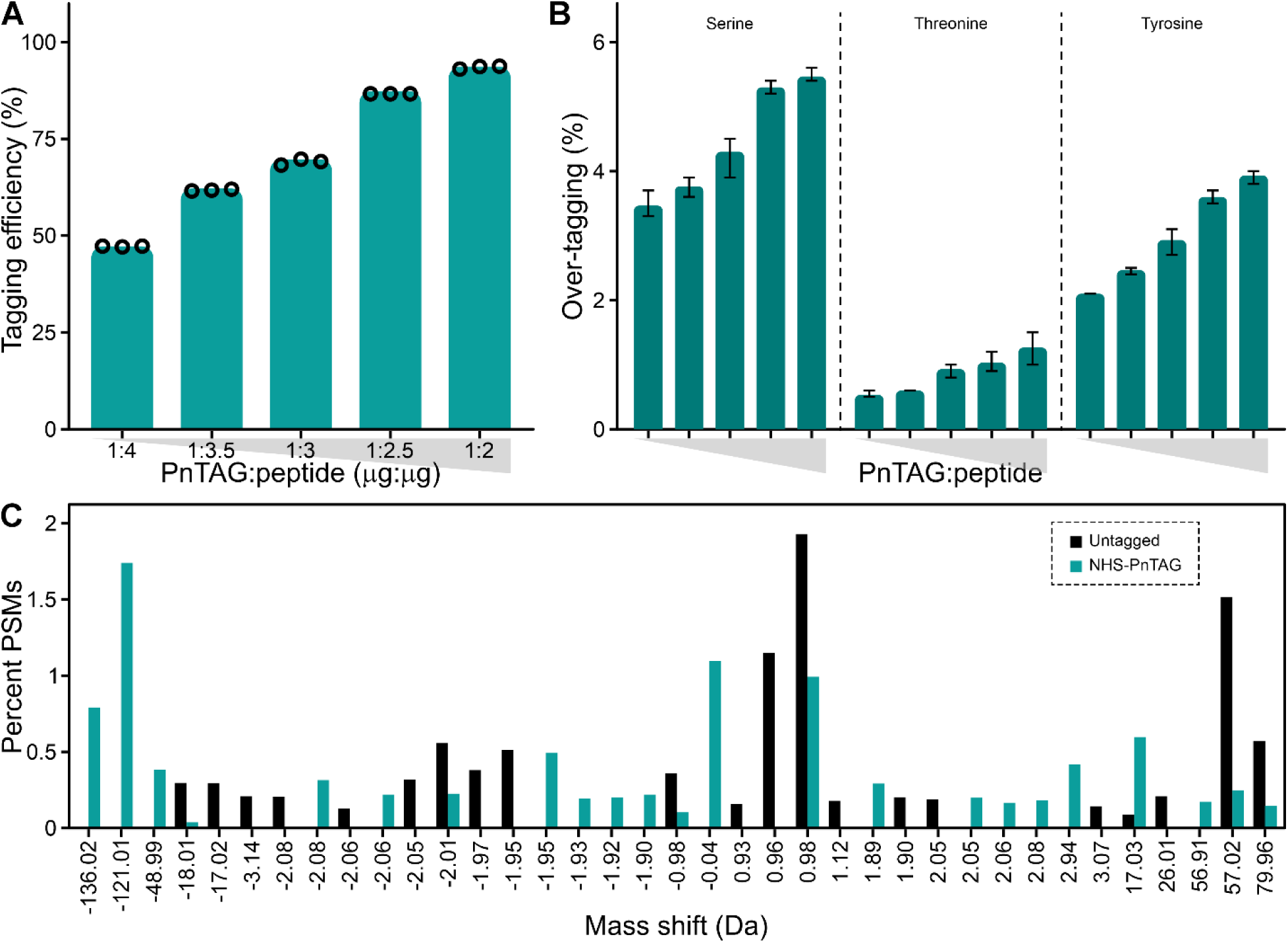
Optimization of internal peptides phosphonate-tagging via NHS ester chemistry. (A) Bar plot showing the proportion of HeLa peptides (by intensity) correctly tagged at lysines and peptide N-termini across different amounts of NHS-PnTAG reagent. (B) Bar plot of the percentage of peptide intensity attributable to over-tagged peptides from the analysis displayed in (A). For all plots, bars are drawn at the mean value (n = 3), dots indicate individual replicates, and error bars indicate the minimum and maximum values. (C) Distribution of the top 15 modifications within each sample identified by MSFragger via the open search strategy and summarized with PTM-Shepherd, with and without phosphonate tagging. Identifications corresponding to unmodified peptides, isotopic mass shifts, or the PnTAG mass shifts were removed from the analysis.

### TMT-SIMPATICO Performance in Identifying Proteolytic Events in Apoptosis

After establishing a streamlined protocol for primary amine blocking and internal peptides tagging, we assessed the ability of SIMPATICO to identify proteolysis-generated termini (Figure 3A). To achieve this, we induced caspase-mediated apoptosis in Jurkat T-cells by treating them with the pan-kinase inhibitor Staurosporine (STS) for six hours. The experiment included five replicates of STS-treated samples and five replicates of DMSO-treated controls (Mock). Cells were lysed in SDC buffer, and primary amines were labeled with TMT10. The combined proteomes underwent direct overnight trypsin digestion and subsequent desalting. Internal peptides were tagged with phosphonate as described above, followed by another step of desalting, HpH RP fractionation, and automated IMAC-based depletion.

**Figure 3.**
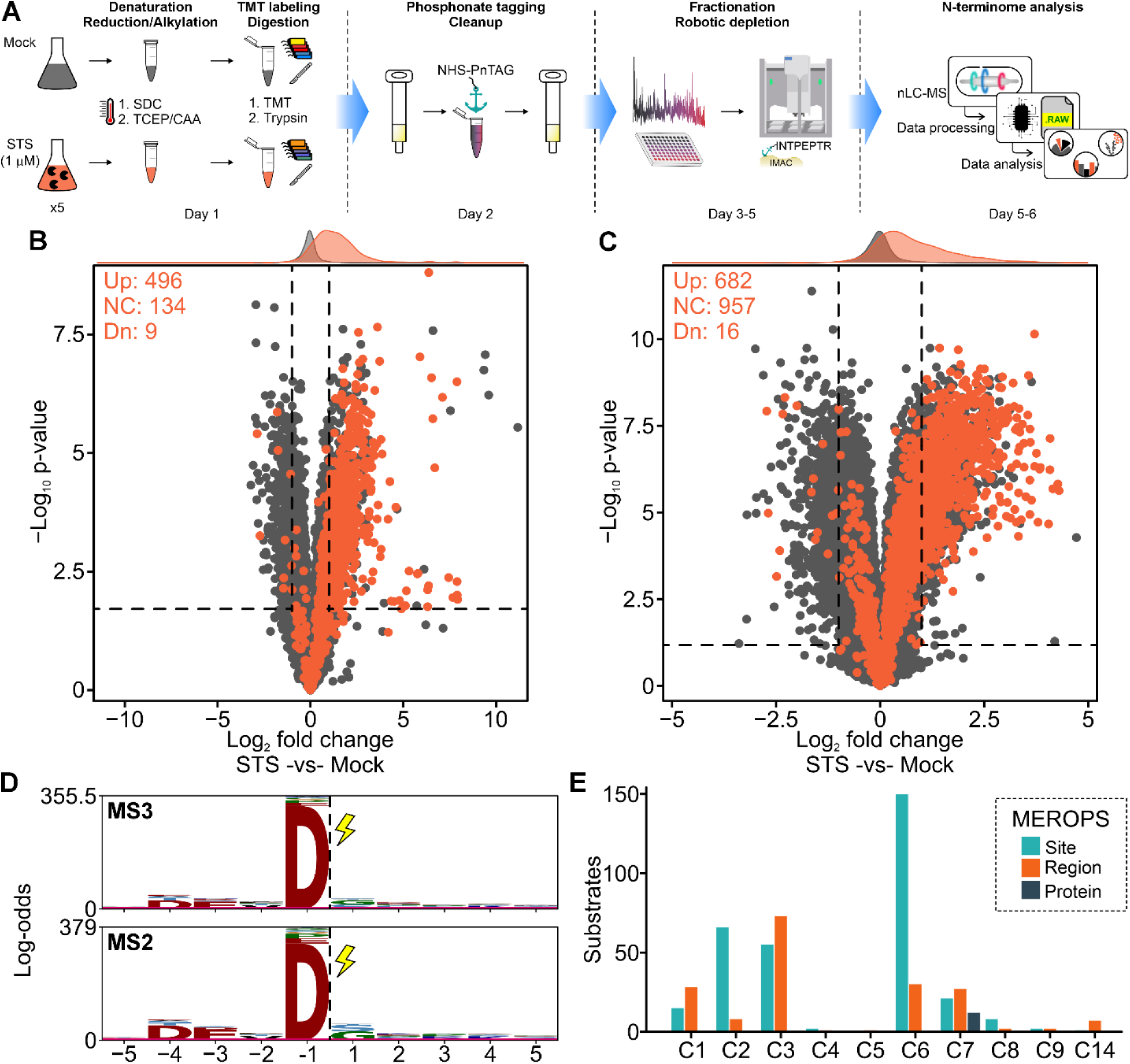
Protocol for streamlined identification of N-terminal peptides. (A) Experimental stages and timing for the TMT-SIMPATICO workflow for the identification of caspase substrates: Protein samples are denatured, reduced, and alkylated. 200 μg of proteins per sample are directly labeled with 600 μg TMT and afterward combined and digested with trypsin (day 1). The resulting peptides are desalted via a C18 cartridge before and after internal peptide tagging with NHS-PnTAG (day 2). The peptide mixture is fractionated via basic pH reverse-phase chromatography, and n-terminal peptides are recovered by negative selection using an automated Fe(III)-IMAC workflow to deplete PnTAG-peptides (days 3-5). N-terminal peptides and cleavage events are identified and quantified via nLC-MS/MS (days 5-6). (B) Volcano plot (Welch’s t-test) showing the N-termini enriched in STS-treated versus Mock-treated Jurkat cells. N-termini generated by cleavage of C-terminal to aspartic residues are highlighted in orange. The *x-axis* shows log2 fold change, while the *y-axis* shows -log10 p-value adjusted for multiple hypothesis testing using the Permutation-based FDR procedure. Dashed lines are at 5% FDR and log2 fold change of +/- 1. Results are from five biological replicates for each condition. For aspartic acid cleavages, the number of upregulated (Up), non-changing (NC), and downregulated (Dn) events is indicated. (C) Same as in (B) but from the MS2-based acquisition. (D) pLogo plots of the enriched cleavages (> 2-fold STS-vs-Mock, FDR < 5%) in response to the STS treatment. Letters larger than the significance threshold (red line) are deemed significantly enriched in those positions. (E) Bar chart showing the potential upstream caspase protease responsible for the generation of the significantly enriched cleavages C-terminal of Asp upon STS treatment, according to the MEROPS database. We defined three different categories describing the information obtained about the cleavage position: explicit cleavage sites (teal), within (orange), or outside (grey) the protein region reported in the database.

We identified over 22,000 n-terminal peptides from approximately 4,500 proteins across all biological replicates (Supplementary Table S3). Since the number of n-terminal peptides largely depends on the proteome’s degradation state under investigation, we first benchmarked our approach against recently established methods for native protein N-termini identification^24,43,44^. Our analysis resulted in the identification of 379 canonical unmodified protein n-termini, with or without the initiator methionine excision, and 1,001 acetylated protein N-termini, resulting in a total of 1,019 non-redundant native n-termini. Although a direct comparison is not possible due to differences in sample types as well as variations in MS instrumentation, input material, sample preparation, and data processing, the values observed here are consistent with previously reported results, demonstrating the capability of SIMPATICO for robust N-terminomics analysis.

Next, to identify apoptosis-induced cleavage events, we performed two-sample *t*-tests comparing STS-treated and mock-treated cells. Significantly upregulated peptides were stringently selected based on a >2-fold increase and a *q*-value < 0.05. In total, SIMPATICO identified 908 cleavage sites that were significantly enriched upon STS treatment (Figure 3B). Notably, asparaginyl cleavages, characteristic of caspase-mediated proteolysis, were prominently enriched in STS-treated samples with 496 unique sites, of which 490 were non-redundant, consistent with activation of apoptotic pathways. Interestingly, several N-terminal peptides show a negative fold change, appearing less abundant in the STS-treated condition, suggesting these N-termini were lost due to cleavage further downstream toward the C-terminus, although the exact cleavage site cannot be pinpointed.

Analyses of the amino acid composition surrounding the 490 cleavage sites clearly showed a preference for small residues such as serine and glycine c-terminal to the cleaved aspartate, as well as the overrepresentation of the DEVD| consensus motif, a well-documented preference of several caspases^45,46^.

Motivated by the high number of cleavage events detected, we sought to further deepen the coverage of the N-terminome by employing a more rapid acquisition method.

To this end, we reanalyzed the same samples using a faster MS2-based acquisition for TMT quantification^47^. This approach enabled the identification of an additional 30,242 N-terminal peptides, 1,842 native protein N-termini, and 390 STS-induced asparaginyl cleavages (Figures 3C, 3D, and Supplementary Table S4).

To generate a list of known caspase substrates for comparison, we compared the STS-induced asparaginyl cleavages detected with SIMPATICO to the MEROPS database^48^. This analysis showed that 202 (36%) of the identified substrates were previously recognized as either verified caspase substrates with exact cleavage sites match or as potential substrates with cleavage evidence within the relevant sequence region or at the protein level (Figure 3E). As expected, given their reported activation by STS^49,50^, CASP-3 and -6 emerged as the most active caspases. The remaining ∼64% of candidates, on 479 proteins, represented yet unreported novel substrates.

Besides potential caspase substrates, our analysis identified STS-induced cleavages at several residues other than aspartate, possibly the result of proteasomal degradation or additional proteolytic events. Notably, 92 glutamyl cleavages (∼5%) were enriched in the STS-treated sample. Closer inspection of the sequence surrounding the cleavage site revealed the preference for the DEVE motif (Supplementary Figure S1), which has been indicated to be cleaved by apoptotic CASP-3 and -7, suggesting that also these proteolytic events are likely mediated by these executioner caspases during apoptosis. In addition, we identified 16 known substrates of cathepsins, lysosomal proteases for which there is growing evidence of activation, although to an unknown extent, during apoptosis^51^, highlighting the strength of SIMPATICO in capturing proteolytic events with high sensitivity across the N-terminome.

### LFQ-SIMPATICO

Quantitative N-terminomics is currently mostly limited to multiplexed quantification methods such as SILAC, dimethyl labeling, or TMT/iTRAQ^20^. Techniques like COFRADIC and its derivatives, although effective for N-terminal enrichment, often require extensive fractionation, which typically restricts their use to the analysis of only a few samples at a time^52^. While multiplexing enables relative quantification, it inherently limits the number of samples that can be analyzed simultaneously. Scaling beyond the multiplexing capacity requires revising the experimental design to include a common reference sample across multiplexing sets^53^, which could become impractical. To date, high-throughput, label-free N-terminomics remains largely unexplored. To address this gap, we investigated the feasibility of adapting our SIMPATICO workflow into a label-free approach.

One of the primary challenges in this adaptation is the efficient blocking of primary amines directly in the lysate. Although TMT0 can be used for this purpose, its high cost makes it unsuitable for large-scale analyses. NHS-acetate, commonly used in COFRADIC-like protocols, also presents limitations: acetylation of primary amines can suppress peptide charge states^54^, as the acetylated nitrogen can no longer retain a proton, thereby hindering MS-based identification of N-terminal peptides. To overcome these limitations, we explored commercially available NHS-based derivatization reagents that would not lead to charge suppression. Candidates such as DMABA NHS-ester and RapiFluor-MS include a tertiary, protonable amine group^55,56^, but both have poor water solubility. We turned our attention to PEG-based NHS esters, a class of amine-reactive reagents with improved water solubility commonly used in click chemistry. We identified DMA-PEG3- NHS as a promising candidate for primary amine blocking (Figure 4A). This reagent contains a tertiary amine as a protonable site and a polyethylene glycol linker for improved solubility in water. Titration experiments demonstrated that DMA-PEG3-NHS can effectively replace TMT for primary amine blocking (Figure 4B, Supplementary Figure S2A, and Supplementary Table S5). Building on this, we evaluated the feasibility of a label-free version of the SIMPATICO workflow to identify proteolysis-generated N-termini. As in previous experiments, Jurkat cells treated with either STS or DMSO were lysed in SDC buffer, and primary amines were blocked using DMA-PEG3-NHS directly in the lysate. Apon tryptic digestion and desalting, internal peptides were selectively tagged with a phosphonate group and removed using zirconia-based SPE. The flow-through from this step was further purified with SDB-RPS, leveraging the dual-mode properties of the sorbent to wash away any undeleted internal peptides from the previous step, which predominantly appear as +1 charge state species due to the acylation of the peptide n- terminus.

**Figure 4.**
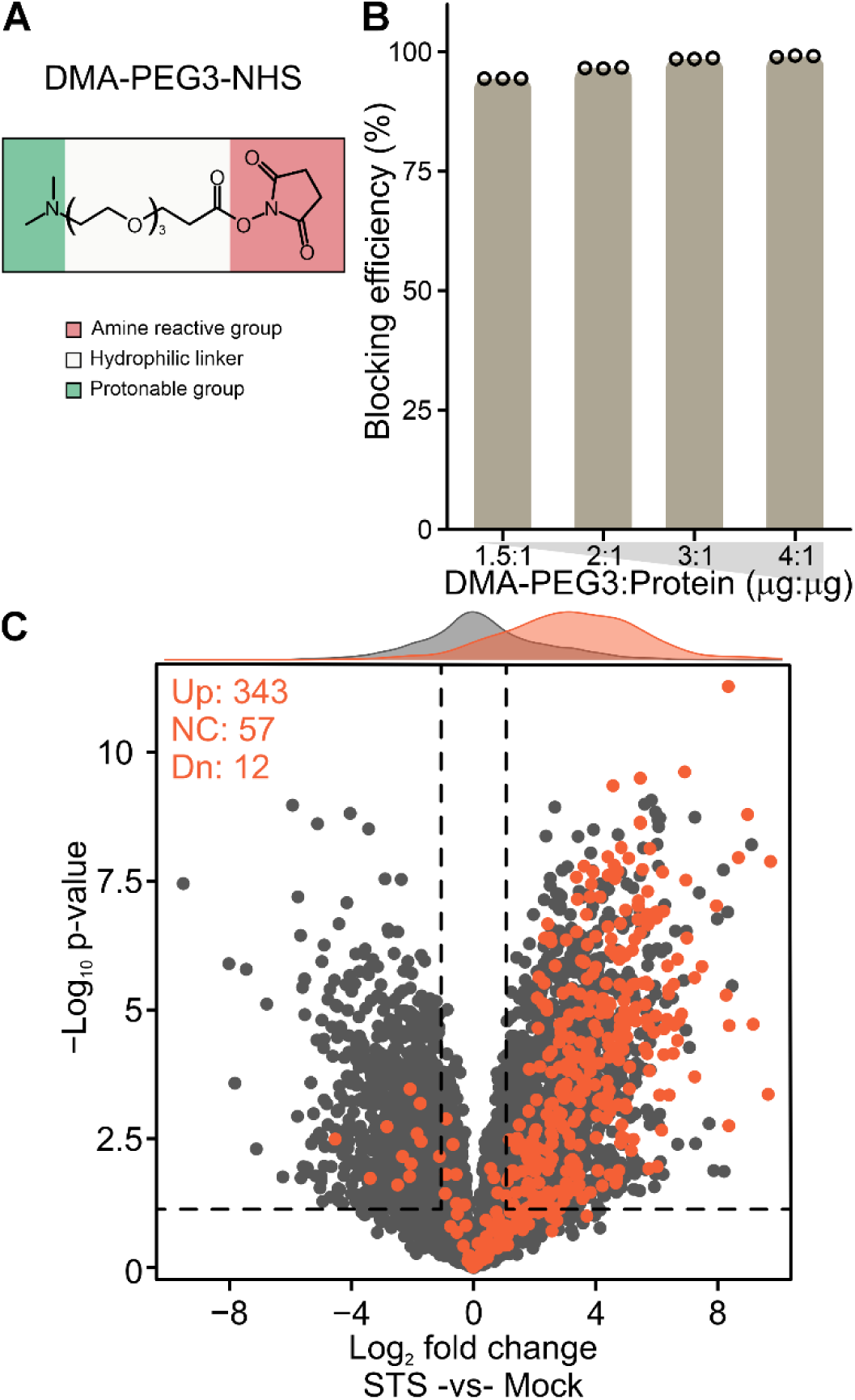
N-terminomics via label-free quantification. (A) Chemical structures of DMA- PEG3-NHS. (B) Bar plot showing the proportion of HeLa peptides (by intensity) correctly blocked at lysines and protein N-termini (blocking efficiency) across different amounts of DMA-PEG3-NHS reagent. Bars are drawn at the mean value (n = 3), and dots indicate individual replicates. (C) Same as in Figure 3B, but from the label-free experiment. Dashed lines are at 5% FDR and log2 fold change of +/- 1. Results are from five biological replicates for each condition.

The enriched N-terminal peptide samples were analyzed via label-free nLC-MS/MS. This analysis successfully identified 5,598 n-terminal peptides, of which 412 were the results of cleavage C-terminal to aspartic residues (Supplementary Table S6). The lower number of detected N-terminal peptides does not come as a surprise. There are several plausible explanations for these observations. In the label-free experiments, no peptide fractionation was performed, meaning that sample complexity remained high and limited the depth of n-terminome coverage. Consequently, in the absence of multiplexing, samples were analyzed individually rather than pooled. This prevented cumulative signal enhancement, so the label-free approach likely captured only the most abundant N-terminal peptides (Supplementary Figure S2B). Furthermore, peptide identifications were not rescored using MSBooster, as the dimethylaminoPEG3ylation is not currently supported for rescoring by predictive models, which leaves room for further improvement in peptide identification. Future efforts could include the use of mild peptide fractionation prior to MS analysis or the adoption of data-independent acquisition (DIA), especially given that recent advances in transfer learning have enabled predictive models to generalize DIA searches to previously unseen modifications^57^.

Nevertheless, 269 aspartyl cleavages (∼65%) were also detected in the multiplexed version of the workflow, and a clear enrichment of aspartyl cleavage events upon STS treatment was also observed (Figure 4C and Supplementary Figure S2C), supporting the effectiveness of the label-free approach for capturing proteolytic signatures.

In summary, we have developed a streamlined sample preparation workflow that minimizes processing steps while enabling efficient and comprehensive N-terminome analysis. By combining a highly reactive NHS-ester reagent with metal coordination-based depletion, this approach allows for robust and selective capture of both canonical and neo-protein N-termini. In its label-free implementation, the workflow can easily be scaled to accommodate up to 96 samples in parallel, making it well-suited for high-throughput studies. We anticipate that this strategy will advance the study of proteolytic processes and offer deeper biological insights, beyond caspase substrate identification.

## ACKNOWLEDGMENTS

This work was supported by the German Research Foundation (DFG) (TRR237 (A07), TRR179 (TP11), TRR353 (B04)). Bavarian State Ministry of Education and Research (BayVFP 2024-2027; P3M) and the Danish National Research Foundation (DNRF 164; CiViA).

The Orbitrap Eclipse and Orbitrap Exploris 480 mass spectrometers were funded in part by the German Research Foundation (INST 95/1650-1 FUGG and INST 95/1649-1 FUGG).

## AUTHOR CONTRIBUTIONS

P.G. performed proteomic experiments, analyzed and interpreted the data; A.F. performed experiments; A.P. supervised the study and acquired funding; P.G. conceptualized the project and wrote the manuscript. All authors edited the manuscript.

## DATA AVAILABILITY

The mass spectrometry proteomics data have been deposited in the ProteomeXchange Consortium via the PRIDE partner repository ^58^ with the dataset identifier PXD064097.

## SUPPLEMENTARY INFORMATION

**Supplementary Figure S1.**
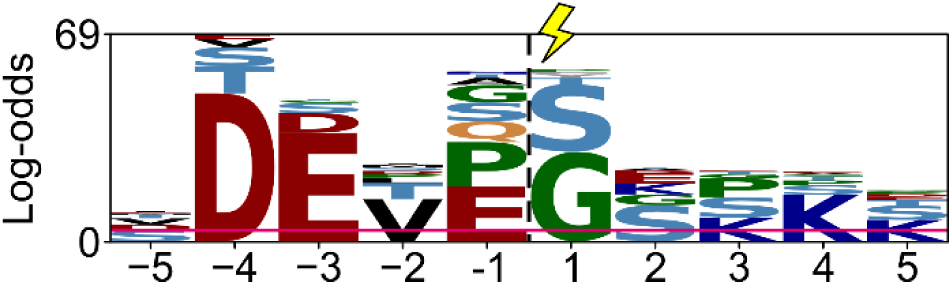
Zoom-in of the pLogo plot shown in Figure 3D (MS3 and MS2 combined data).

**Supplementary Figure S2.**
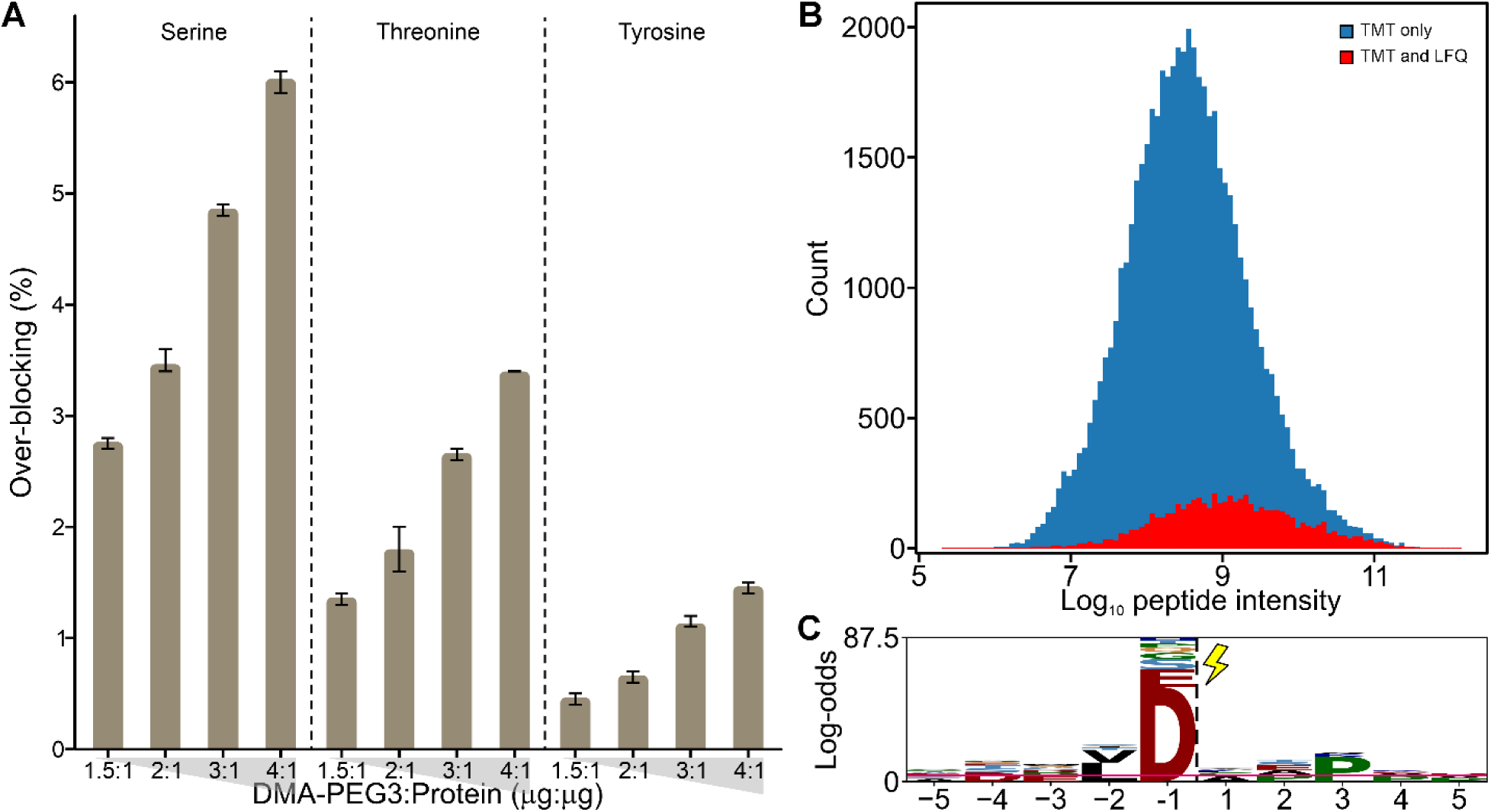
(A) Bar plot of the percentage of peptide intensity attributable to over-blocked peptides across different amounts of DMA-PEG3-NHS reagent, from the analysis displayed in Figure 4B. Bars are drawn at the mean value (n = 3); error bars indicate the minimum and maximum values. (B) Histogram of log n-terminal peptide quantified in the TMT-SIMPATICO experiment from the MS2-based quantification. Red bars indicate N-terminal peptides that were also detected in the LFQ experiment. These peptides are predominantly found at higher intensity values within the distribution. (C) pLogo plot of the enriched cleavages (> 2-fold STS-vs-Mock, FDR < 5%) in response to the STS treatment from the LFQ experiment.

**Supplementary Table S1.** TMT protein titration experiments.

**Supplementary Table S2.** NHS-PnTAG peptide titration experiments.

**Supplementary Table S3.** N-terminal peptides identification by TMT-SIMPATICO with an MS3-based quantification method.

**Supplementary Table S4.** N-terminal peptides identification by TMT-SIMPATICO with an MS2-based quantification method.

**Supplementary Table S5.** DMA-PEG3-NHS protein titration experiments.

**Supplementary Table S6.** N-terminal peptides identification by LFQ-SIMPATICO.

## REFERENCES

(1) Hinkson, I. V.; Elias, J. E. The Dynamic State of Protein Turnover: It’s about Time. Trends Cell Biol. 2011, 21 (5), 293–303. 10.1016/j.tcb.2011.02.002.

(2) Strasser, A.; O’Connor, L.; Dixit, V. M. Apoptosis Signaling. Annu. Rev. Biochem. 2000, 69, 217–245. 10.1146/annurev.biochem.69.1.217.

(3) Sladky, V. C.; Villunger, A. Uncovering the PIDDosome and Caspase-2 as Regulators of Organogenesis and Cellular Differentiation. Cell Death Differ. 2020, 27 (7), 2037–2047. 10.1038/s41418-020-0556-6.

(4) Rai, M.; Curley, M.; Coleman, Z.; Demontis, F. Contribution of Proteases to the Hallmarks of Aging and to Age-Related Neurodegeneration. Aging Cell 2022, 21 (5), e13603. 10.1111/acel.13603.

(5) Bunnett, N. W. Protease-Activated Receptors: How Proteases Signal to Cells to Cause Inflammation and Pain. Semin. Thromb. Hemost. 2006, 32 *Suppl 1*, 39–48. 10.1055/s-2006-939553.

(6) Lloyd, R. E. Enterovirus Control of Translation and RNA Granule Stress Responses. Viruses 2016, 8 (4), 93. 10.3390/v8040093.

(7) Borges, P. H. O.; Ferreira, S. B.; Silva, F. P. Recent Advances on Targeting Proteases for Antiviral Development. Viruses 2024, 16 (3), 366. 10.3390/v16030366.

(8) Tözsér, J.; Oroszlan, S. Proteolytic Events of HIV-1 Replication as Targets for Therapeutic Intervention. Curr. Pharm. Des. 2003, 9 (22), 1803–1815. 10.2174/1381612033454478.

(9) Mason, S. D.; Joyce, J. A. Proteolytic Networks in Cancer. Trends Cell Biol. 2011, 21 (4), 10.1016/j.tcb.2010.12.002.

(10) Klingler, D.; Hardt, M. Profiling Protease Activities by Dynamic Proteomics Workflows. Proteomics 2012, 12 (4–5), 587–596. 10.1002/pmic.201100399.

(11) McShane, E.; Sin, C.; Zauber, H.; Wells, J. N.; Donnelly, N.; Wang, X.; Hou, J.; Chen, W.; Storchova, Z.; Marsh, J. A.; Valleriani, A.; Selbach, M. Kinetic Analysis of Protein Stability Reveals Age-Dependent Degradation. Cell 2016, 167 (3), 803–815.e21. 10.1016/j.cell.2016.09.015.

(12) Zecha, J.; Meng, C.; Zolg, D. P.; Samaras, P.; Wilhelm, M.; Kuster, B. Peptide Level Turnover Measurements Enable the Study of Proteoform Dynamics. Mol. Cell. Proteomics MCP 2018, 17 (5), 974–992. 10.1074/mcp.RA118.000583.

(13) Julien, O.; Zhuang, M.; Wiita, A. P.; O’Donoghue, A. J.; Knudsen, G. M.; Craik, C. S.; Wells, J. A. Quantitative MS-Based Enzymology of Caspases Reveals Distinct Protein Substrate Specificities, Hierarchies, and Cellular Roles. Proc. Natl. Acad. Sci. U. S. A. 2016, 113 (14), E2001–2010. 10.1073/pnas.1524900113.

(14) O’Donoghue, A. J.; Eroy-Reveles, A. A.; Knudsen, G. M.; Ingram, J.; Zhou, M.; Statnekov, J. B.; Greninger, A. L.; Hostetter, D. R.; Qu, G.; Maltby, D. A.; Anderson, M. O.; Derisi, J. L.; McKerrow, J. H.; Burlingame, A. L.; Craik, C. S. Global Identification of Peptidase Specificity by Multiplex Substrate Profiling. Nat. Methods 2012, 9 (11), 1095–1100. 10.1038/nmeth.2182.

(15) Anania, V. G.; Yu, K.; Gnad, F.; Pferdehirt, R. R.; Li, H.; Ma, T. P.; Jeon, D.; Fortelny, N.; Forrest, W.; Ashkenazi, A.; Overall, C. M.; Lill, J. R. Uncovering a Dual Regulatory Role for Caspases During Endoplasmic Reticulum Stress-Induced Cell Death. Mol. Cell. Proteomics MCP 2016, 15 (7), 2293–2307. 10.1074/mcp.M115.055376.

(16) Dix, M. M.; Simon, G. M.; Cravatt, B. F. Global Mapping of the Topography and Magnitude of Proteolytic Events in Apoptosis. Cell 2008, 134 (4), 679–691. 10.1016/j.cell.2008.06.038.

(17) Dix, M. M.; Simon, G. M.; Wang, C.; Okerberg, E.; Patricelli, M. P.; Cravatt, B. F. Functional Interplay between Caspase Cleavage and Phosphorylation Sculpts the Apoptotic Proteome. Cell 2012, 150 (2), 426–440. 10.1016/j.cell.2012.05.040.

(18) Stoehr, G.; Schaab, C.; Graumann, J.; Mann, M. A SILAC-Based Approach Identifies Substrates of Caspase-Dependent Cleavage upon TRAIL-Induced Apoptosis. Mol. Cell. Proteomics MCP 2013, 12 (5), 1436–1450. 10.1074/mcp.M112.024679.

(19) McDonald, L.; Robertson, D. H. L.; Hurst, J. L.; Beynon, R. J. Positional Proteomics: Selective Recovery and Analysis of N-Terminal Proteolytic Peptides. Nat. Methods 2005, 2 (12), 955–957. 10.1038/nmeth811.

(20) Luo, S. Y.; Araya, L. E.; Julien, O. Protease Substrate Identification Using N-Terminomics. ACS Chem. Biol. 2019, 14 (11), 2361–2371. 10.1021/acschembio.9b00398.

(21) Kleifeld, O.; Doucet, A.; auf dem Keller, U.; Prudova, A.; Schilling, O.; Kainthan, R. K.; Starr, A. E.; Foster, L. J.; Kizhakkedathu, J. N.; Overall, C. M. Isotopic Labeling of Terminal Amines in Complex Samples Identifies Protein N-Termini and Protease Cleavage Products. Nat. Biotechnol. 2010, 28 (3), 281–288. 10.1038/nbt.1611.

(22) Weng, S. S. H.; Demir, F.; Ergin, E. K.; Dirnberger, S.; Uzozie, A.; Tuscher, D.; Nierves, L.; Tsui, J.; Huesgen, P. F.; Lange, P. F. Sensitive Determination of Proteolytic Proteoforms in Limited Microscale Proteome Samples. Mol. Cell. Proteomics MCP 2019, 18 (11), 2335–2347. 10.1074/mcp.TIR119.001560.

(23) Kalogeropoulos, K.; Bundgaard, L.; Auf dem Keller, U. Sensitive and High-Throughput Exploration of Protein N-Termini by TMT-TAILS N-Terminomics. Methods Mol. Biol. Clifton NJ 2023, 2718, 111–135. 10.1007/978-1-0716-3457-8_7.

(24) Demir, F.; Perrar, A.; Mantz, M.; Huesgen, P. F. Sensitive Plant N-Terminome Profiling with HUNTER. Methods Mol. Biol. Clifton NJ 2022, 2447, 139–158. 10.1007/978-1-0716-2079-3_12.

(25) Baghalabadi, V.; Doucette, A. A. Mass Spectrometry Profiling of Low Molecular Weight Proteins and Peptides Isolated by Acetone Precipitation. Anal. Chim. Acta 2020, 1138, 38–48. 10.1016/j.aca.2020.08.057.

(26) Ahrens, C. H.; Wade, J. T.; Champion, M. M.; Langer, J. D. A Practical Guide to Small Protein Discovery and Characterization Using Mass Spectrometry. J. Bacteriol. 2022, 204 (1), e0035321. 10.1128/JB.00353-21.

(27) MacDonald, B. T.; Keshishian, H.; Mundorff, C. C.; Arduini, A.; Lai, D.; Bendinelli, K.; Popp, N. R.; Bhandary, B.; Clauser, K. R.; Specht, H.; Elowe, N. H.; Laprise, D.; Xing, Y.; Kaushik, V. K.; Carr, S. A.; Ellinor, P. T. TAILS Identifies Candidate Substrates and Biomarkers of ADAMTS7, a Therapeutic Protease Target in Coronary Artery Disease. Mol. Cell. Proteomics MCP 2022, 21 (4), 100223. 10.1016/j.mcpro.2022.100223.

(28) Kalogeropoulos, K.; Moldt Haack, A.; Madzharova, E.; Di Lorenzo, A.; Hanna, R.; Schoof, E. M.; Auf dem Keller, U. CLIPPER 2.0: Peptide-Level Annotation and Data Analysis for Positional Proteomics. Mol. Cell. Proteomics MCP 2024, 23 (6), 100781. 10.1016/j.mcpro.2024.100781.

(29) Mommen, G. P. M.; van de Waterbeemd, B.; Meiring, H. D.; Kersten, G.; Heck, A. J. R.; de Jong, A. P. J. M. Unbiased Selective Isolation of Protein N-Terminal Peptides from Complex Proteome Samples Using Phospho Tagging (PTAG) and TiO(2)- Based Depletion. Mol. Cell. Proteomics MCP 2012, 11 (9), 832–842. 10.1074/mcp.O112.018283.

(30) Zecha, J.; Satpathy, S.; Kanashova, T.; Avanessian, S. C.; Kane, M. H.; Clauser, K. R.; Mertins, P.; Carr, S. A.; Kuster, B. TMT Labeling for the Masses: A Robust and Cost-Efficient, In-Solution Labeling Approach. Mol. Cell. Proteomics MCP 2019, 18 (7), 1468–1478. 10.1074/mcp.TIR119.001385.

(31) Post, H.; Penning, R.; Fitzpatrick, M. A.; Garrigues, L. B.; Wu, W.; MacGillavry, H. D.; Hoogenraad, C. C.; Heck, A. J. R.; Altelaar, A. F. M. Robust, Sensitive, and Automated Phosphopeptide Enrichment Optimized for Low Sample Amounts Applied to Primary Hippocampal Neurons. J. Proteome Res. 2017, 16 (2), 728–737. 10.1021/acs.jproteome.6b00753.

(32) Kong, A. T.; Leprevost, F. V.; Avtonomov, D. M.; Mellacheruvu, D.; Nesvizhskii, A. I. MSFragger: Ultrafast and Comprehensive Peptide Identification in Mass Spectrometry–Based Proteomics. Nat. Methods 2017, 14 (5), 513–520. 10.1038/nmeth.4256.

(33) Yang, K. L.; Yu, F.; Teo, G. C.; Li, K.; Demichev, V.; Ralser, M.; Nesvizhskii, A. I. MSBooster: Improving Peptide Identification Rates Using Deep Learning-Based Features. Nat. Commun. 2023, 14 (1), 4539. 10.1038/s41467-023-40129-9.

(34) Lautenbacher, L.; Yang, K. L.; Kockmann, T.; Panse, C.; Chambers, M.; Kahl, E.; Yu, F.; Gabriel, W.; Bold, D.; Schmidt, T.; Li, K.; MacLean, B.; Nesvizhskii, A. I.; Wilhelm, M. Koina: Democratizing Machine Learning for Proteomics Research. bioRxiv June 3, 2024, p 2024.06.01.596953. 10.1101/2024.06.01.596953.

(35) Käll, L.; Canterbury, J. D.; Weston, J.; Noble, W. S.; MacCoss, M. J. Semi-Supervised Learning for Peptide Identification from Shotgun Proteomics Datasets. Nat. Methods 2007, 4 (11), 923–925. 10.1038/nmeth1113.

(36) da Veiga Leprevost, F.; Haynes, S. E.; Avtonomov, D. M.; Chang, H.-Y.; Shanmugam, A. K.; Mellacheruvu, D.; Kong, A. T.; Nesvizhskii, A. I. Philosopher: A Versatile Toolkit for Shotgun Proteomics Data Analysis. Nat. Methods 2020, 17 (9), 869–870. 10.1038/s41592-020-0912-y.

(37) Tyanova, S.; Cox, J. Perseus: A Bioinformatics Platform for Integrative Analysis of Proteomics Data in Cancer Research. Methods Mol. Biol. Clifton NJ 2018, 1711, 133–148. 10.1007/978-1-4939-7493-1_7.

(38) León, I. R.; Schwämmle, V.; Jensen, O. N.; Sprenger, R. R. Quantitative Assessment of In-Solution Digestion Efficiency Identifies Optimal Protocols for Unbiased Protein Analysis*. Mol. Cell. Proteomics 2013, 12 (10), 2992–3005. 10.1074/mcp.M112.025585.

(39) Chen, L.; Shan, Y.; Weng, Y.; Sui, Z.; Zhang, X.; Liang, Z.; Zhang, L.; Zhang, Y. Hydrophobic Tagging-Assisted N-Termini Enrichment for In-Depth N-Terminome Analysis. Anal. Chem. 2016, 88 (17), 8390–8395. 10.1021/acs.analchem.6b02453.

(40) Steigenberger, B.; Pieters, R. J.; Heck, A. J. R.; Scheltema, R. A. PhoX: An IMAC-Enrichable Cross-Linking Reagent. ACS Cent. Sci. 2019, 5 (9), 1514–1522. 10.1021/acscentsci.9b00416.

(41) Kleinpenning, F.; Steigenberger, B.; Wu, W.; Heck, A. J. R. Fishing for Newly Synthesized Proteins with Phosphonate-Handles. Nat. Commun. 2020, 11 (1), 3244. 10.1038/s41467-020-17010-0.

(42) Liu, X.; Rossio, V.; Gygi, S. P.; Paulo, J. A. Enriching Cysteine-Containing Peptides Using a Sulfhydryl-Reactive Alkylating Reagent with a Phosphonic Acid Group and Immobilized Metal Affinity Chromatography. J. Proteome Res. 2023, 22 (4), 1270–1279. 10.1021/acs.jproteome.2c00806.

(43) Venne, A. S.; Solari, F. A.; Faden, F.; Paretti, T.; Dissmeyer, N.; Zahedi, R. P. An Improved Workflow for Quantitative N-Terminal Charge-Based Fractional Diagonal Chromatography (ChaFRADIC) to Study Proteolytic Events in Arabidopsis Thaliana. Proteomics 2015, 15 (14), 2458–2469. 10.1002/pmic.201500014.

(44) Yeom, J.; Ju, S.; Choi, Y.; Paek, E.; Lee, C. Comprehensive Analysis of Human Protein N-Termini Enables Assessment of Various Protein Forms. Sci. Rep. 2017, 7 (1), 6599. 10.1038/s41598-017-06314-9.

(45) Julien, O.; Wells, J. A. Caspases and Their Substrates. Cell Death Differ. 2017, 24 (8), 1380–1389. 10.1038/cdd.2017.44.

(46) Seaman, J. E.; Julien, O.; Lee, P. S.; Rettenmaier, T. J.; Thomsen, N. D.; Wells, J. A. Cacidases: Caspases Can Cleave after Aspartate, Glutamate and Phosphoserine Residues. Cell Death Differ. 2016, 23 (10), 1717–1726. 10.1038/cdd.2016.62.

(47) Hogrebe, A.; von Stechow, L.; Bekker-Jensen, D. B.; Weinert, B. T.; Kelstrup, C. D.; Olsen, J. V. Benchmarking Common Quantification Strategies for Large-Scale Phosphoproteomics. Nat. Commun. 2018, 9 (1), 1045. 10.1038/s41467-018-03309-6.

(48) Rawlings, N. D.; Barrett, A. J.; Thomas, P. D.; Huang, X.; Bateman, A.; Finn, R. D. The MEROPS Database of Proteolytic Enzymes, Their Substrates and Inhibitors in 2017 and a Comparison with Peptidases in the PANTHER Database. Nucleic Acids Res. 2018, 46 (D1), D624–D632. 10.1093/nar/gkx1134.

(49) Warby, S. C.; Doty, C. N.; Graham, R. K.; Carroll, J. B.; Yang, Y.-Z.; Singaraja, R. R.; Overall, C. M.; Hayden, M. R. Activated Caspase-6 and Caspase-6-Cleaved Fragments of Huntingtin Specifically Colocalize in the Nucleus. Hum. Mol. Genet. 2008, 17 (15), 2390–2404. 10.1093/hmg/ddn139.

(50) Thuret, G.; Chiquet, C.; Herrag, S.; Dumollard, J.-M.; Boudard, D.; Bednarz, J.; Campos, L.; Gain, P. Mechanisms of Staurosporine Induced Apoptosis in a Human Corneal Endothelial Cell Line. Br. J. Ophthalmol. 2003, 87 (3), 346–352.

(51) Turk, B.; Stoka, V.; Rozman-Pungercar, J.; Cirman, T.; Droga-Mazovec, G.; Oresić, K.; Turk, V. Apoptotic Pathways: Involvement of Lysosomal Proteases. Biol. Chem. 2002, 383 (7–8), 1035–1044. 10.1515/BC.2002.112.

(52) Staes, A.; Van Damme, P.; Timmerman, E.; Ruttens, B.; Stes, E.; Gevaert, K.; Impens, F. Protease Substrate Profiling by N-Terminal COFRADIC. Methods Mol. Biol. Clifton NJ 2017, 1574, 51–76. 10.1007/978-1-4939-6850-3_5.

(53) Plubell, D. L.; Wilmarth, P. A.; Zhao, Y.; Fenton, A. M.; Minnier, J.; Reddy, A. P.; Klimek, J.; Yang, X.; David, L. L.; Pamir, N. Extended Multiplexing of Tandem Mass Tags (TMT) Labeling Reveals Age and High Fat Diet Specific Proteome Changes in Mouse Epididymal Adipose Tissue. Mol. Cell. Proteomics MCP 2017, 16 (5), 873. 10.1074/mcp.M116.065524.

(54) Zolg, D. P.; Wilhelm, M.; Schmidt, T.; Médard, G.; Zerweck, J.; Knaute, T.; Wenschuh, H.; Reimer, U.; Schnatbaum, K.; Kuster, B. ProteomeTools: Systematic Characterization of 21 Post-Translational Protein Modifications by Liquid Chromatography Tandem Mass Spectrometry (LC-MS/MS) Using Synthetic Peptides. Mol. Cell. Proteomics MCP 2018, 17 (9), 1850–1863. 10.1074/mcp.TIR118.000783.

(55) Berry, K. A. Z.; Turner, W. W.; VanNieuwenhze, M. S.; Murphy, R. C. Stable Isotope Labeled 4-(Dimethylamino)Benzoic Acid Derivatives of Glycerophosphoethanolamine Lipids. Anal. Chem. 2009, 81 (16), 6633–6640. 10.1021/ac900583a.

(56) Lauber, M. A.; Yu, Y.-Q.; Brousmiche, D. W.; Hua, Z.; Koza, S. M.; Magnelli, P.; Guthrie, E.; Taron, C. H.; Fountain, K. J. Rapid Preparation of Released N-Glycans for HILIC Analysis Using a Labeling Reagent That Facilitates Sensitive Fluorescence and ESI-MS Detection. Anal. Chem. 2015, 87 (10), 5401–5409. 10.1021/acs.analchem.5b00758.

(57) Wallmann, G.; Skowronek, P.; Brennsteiner, V.; Lebedev, M.; Thielert, M.; Steigerwald, S.; Kotb, M.; Heymann, T.; Zhou, X.-X.; Schwörer, M.; Strauss, M. T.; Ammar, C.; Willems, S.; Zeng, W.-F.; Mann, M. AlphaDIA Enables End-to-End Transfer Learning for Feature-Free Proteomics. bioRxiv June 2, 2024, p 2024.05.28.596182. 10.1101/2024.05.28.596182.

(58) Perez-Riverol, Y.; Bandla, C.; Kundu, D. J.; Kamatchinathan, S.; Bai, J.; Hewapathirana, S.; John, N. S.; Prakash, A.; Walzer, M.; Wang, S.; Vizcaíno, J. A. The PRIDE Database at 20 Years: 2025 Update. Nucleic Acids Res. 2025, 53 (D1), D543–D553. 10.1093/nar/gkae1011.

